# Subnanometer-resolution structure determination *in situ* by a hybrid subtomogram averaging – single particle cryoEM – workflow

**DOI:** 10.1101/2020.04.24.057299

**Authors:** Ricardo Sanchez, Yingyi Zhang, Wenbo Chen, Lea Dietrich, Misha Kudryashev

**Affiliations:** Max Planck Institute for Biophysics, Max-von-Laue Strasse, 3, Frankfurt am Main, Germany; Buchmann Institute for Molecular Life Sciences, Goethe University of Frankfurt am Main, Max-von-Laue Strasse, 15, Frankfurt on Main, Germany; Department of Structural Biology, Max Planck Institute for Biophysics, Max-von-Laue Strasse, 3, Frankfurt am Main, Germany

## Abstract

Cryo electron tomography (cryo-ET) combined with subtomogram averaging (StA) enables structural determination of macromolecules in their native context. A few structures were reported by StA at resolution higher than 4.5 Å, however all of these are from viral structural proteins or vesicle coats. Reaching high resolution for a broader range of samples is uncommon due to beam-induced sample drift, poor signal-to-noise ratio (SNR) of images, challenges in CTF correction, limited number of particles. Here we propose a strategy to address these issues, which consists of a tomographic data collection scheme and a processing workflow. Tilt series are collected with higher electron dose at zero-degree tilt in order to increase SNR. Next, after performing StA conventionally, we extract 2D projections of the particles of interest from the higher SNR images and use the single particle analysis tools to refine the particle alignment and generate a reconstruction. We benchmarked our proposed *hybrid StA (hStA)* workflow and improved the resolution for tobacco mosaic virus from 7.2 to 5.2 Å and the resolution for the ion channel RyR1 in crowded native membranes from 12.9 to 9.1 Å. We demonstrate that *hStA* can improve the resolution obtained by conventional StA and promises to be a useful tool for StA projects aiming at subnanometer resolution or higher.

## INTRODUCTION

Cryo electron microscopy (cryo-EM) is a versatile technique for achieving precise structural knowledge of biological processes over a range of scales. Single particle cryo-EM (cryo-SPA) is capable of yielding high resolution structures of purified proteins up to 1.62 Å (Danev *et al*, 2019), with the average resolution of the maps, deposited in the Electron Microscopy Data Bank (EMDB) in 2019, hitting 5.6 Å. In contrast, the average resolution of the StA structures deposited in 2019 is ∼28 Å, although a few structures of viral proteins have been reported in the range of 3 Å to 4 Å resolution (Dick *et al*, 2020, Turoňová *et al*, 2020, Schur *et al*, 2016, Turoňová *et al*, 2017, Himes & Zhang, 2018). Furthermore, even at subnanometer resolution, StA structures are of great interest as they allow resolving secondary structural elements and provide unique insights into the function of proteins in their native context (Kovtun *et al*, 2018,O’Riley, 2020,Wan *et al*, 2017,Hutchings *et al*, 2018).

The *conventional* StA workflow (Leigh *et al*, 2019) starts by (1) recording dose-fractionated movies at each of the projection angles in a tilt series, (2) aligning each of the motion-corrected tilt images in the series, typically using gold fiducials, (3) estimating the defocus and performing contrast transfer function (CTF) correction on the aligned stacks by phase flipping, and (4) generating tomographic reconstructions using the CTF-corrected aligned stacks. Next, the reconstructed tomograms are used for (5) particle picking and extraction followed by (6) alignment of the particles to a common average with optional classification. Additional steps may include 3D CTF correction (Kunz & Frangakis, 2016,Turoňová *et al*, 2017,Bharat *et al*, 2015), correction for local sample deformation based on gold fiducial alignment (Fernandez *et al*, 2019), and/or constrained refinement of tomographic geometry based on the positions of the particles (Bartesaghi *et al*, 2012,Himes & Zhang, 2018). Conventional StA workflow faces several bottlenecks, which make achieving high resolution challenging:

- Beam-induced specimen movement is not uniform (Brilot *et al*, 2012), and may surpass 10 Å (Campbell *et al*, 2012). As a result, tomograms are not recorded from rigid objects, but rather from ones that are continuously changing as the electron dose accumulates.
- Due to the distribution of the total electron dose of 60-150 e^−^/Å^2^ over 40-60 projections, the precision of defocus determination is lower than in single particle cryo-EM.
- The effective thickness of the sample is increasing during tilting and introducing an additional defocus gradient in the direction of the electron beam. This leads to further difficulties in defocus determination and CTF correction. CTF correction error of 250 nm limits the maximum achievable resolution to ∼10 Å, on a 300 kV microscope (Kudryashev, 2018).
- A large number of particles is required to be collected and processed, in order to achieve high resolution. Recording tomograms at lower magnification provides higher throughput, however comes at a price of data quality (Leigh *et al*, 2019) and anisotropic magnification distortions (Grant & Grigorieff, 2015).

Here we present a workflow which combines the advantages of single particle cryo-EM and StA with a focus on achieving subnanometer resolution. Our workflow consists of recording a higher dose micrograph at zero degree followed by recording the remainder of the tilt series in the standard way, incorporating the “high-dose” image into the tilt series and performing StA using the described above workflow. At the last step the particles are extracted in 2D from the “high-dose” images are locally refined against their average. We refer to the tomograms recorded using this method as hybrid tomograms and the entire workflow as hybrid subtomogram averaging (*hStA*). We first tested *hStA* on tobacco mosaic virus (TMV) recorded at a relatively low magnification of an electron microscope corresponding to 2.2 Å/pixel in counted mode, with an aim to collect a larger particle number. We compared the performance of conventional StA processing of tomograms recorded with the standard dose-symmetric tilt scheme (Hagen *et al*, 2016) against conventional StA of hybrid tomograms. We further evaluated the performance of *hStA* on an ion channel RyR1 preserved in native sarcoplasmic reticulum (SR) membranes isolated from rabbit skeletal muscle. Our results show significant improvement in resolution by using *hStA* compared to the conventional StA.

## RESULTS

### Hybrid StA-SPA (*hStA*) Workflow

We extended the dose-symmetric SerialEM script (Hagen *et al*, 2016) to record a “high dose” movie as the first exposure of the tilt series (script available in the Supplementary note 1). The electron dose for this “high dose” movie is 15 e^−^/Å^2^, with the total dose for the entire tilt series equal to approximately 95 e^−^/Å^2^ (Figure 1A). Anisotropic motion correction is performed on the “high dose” micrograph using MotionCor2 (Zheng *et al*, 2017) with the last frame used as the reference. The movies for the remaining tilts are aligned globally. The micrographs are normalized to have the same mean values; their standard deviations are adjusted according to the electron dose applied for each micrograph with the “high dose” image resulting in lower standard deviation. We assumed the following empirical relationship: (σ_HD_)^2^/ (σ_LD_)^2^ = e^−^_LD_ / e^−^ HD. For our data collection scheme, with e^−^_HD_ = 15 e^−^/Å^2^ and e^−^_LD_ = 2 e^−^/Å^2^, the ratio of standard deviations is σ_HD_ = 0.37 * σ_LD_. The tilt series are then aligned using gold beads and tomograms are reconstructed conventionally using Imod (Kremer *et al*, 1996), the particles are identified in the tomograms and StA is performed using Dynamo (Castaño-Díez *et al*, 2012) (Figure 1B). Based on the final StA alignment, the particles are located in the original “high-dose” image, and extracted in 2D. The rotations needed to align these 2D particles to the average are calculated as a combination of the geometrical transforms from the StA workflow (Figure 1B). In our implementation, these rotations are stored in a widely used STAR-formatted file in the ZYZ Euler angles representation. The defocus difference relative to the center of the tomogram is known for each particle, based on the Z-heights of the particle in the tomogram. The updated defocus values are also stored in the STAR file. We refer to the reconstruction resulting from the “high dose” micrographs as a *hybrid map*. The positions and rotations of the particles are further refined using this hybrid map as a reference, with the resulting reconstruction referred to as the *refined map*. During this refinement step, using Relion 3.0 (Zivanov *et al*, 2018), only small local searches are performed to the translations and rotations of each particle. The processing script and the usage instructions can be found in Supplementary note 2.

**Figure 1.**
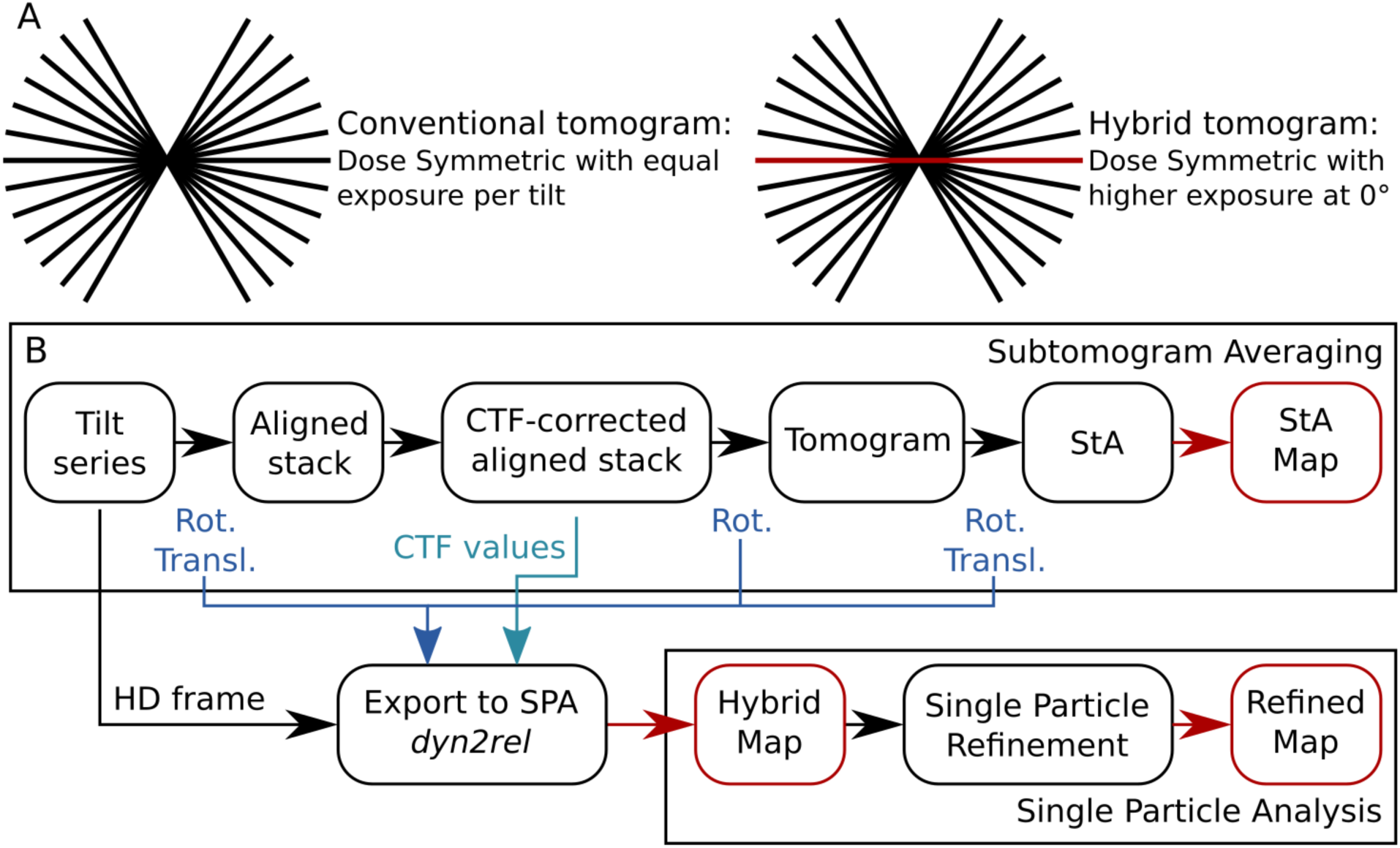
Scheme of the Hybrid StA-SPA workflow. A) A schematic depicting the distribution of dose in the dose-symmetric scheme (Hagen *et al*, 2016) with uniform dose distribution (left) and with an increased exposure for the untilted image shown in red (right panel). B) Data flow diagram for conventional and hybrid StA, black boxes correspond to the intermediate steps, red boxes indicate the output maps, blue arrows indicate geometric transforms performed between the processing steps and the CTF information which are exported for hybrid StA processing.

### Resolution gain with hStA

In order to assess the benefits of our approach experimentally, we first used the highly stable and helically symmetric tobacco mosaic virus (TMV) as a test sample. We acquired tomograms on a Titan Krios (Thermo Fisher Scientific) equipped with a K2 camera (Gatan) and energy filter at an intermediate nominal magnification of 64 000x (2.2 Å/pixel in counting mode) with the aim of imaging larger fields of view and therefore more particles. We recorded small datasets of conventional dose-symmetric tomograms (4,570 particles) and “hybrid tomograms” (4,190 particles), with an equivalent total electron dose of approximately 95 e^−^/Å^2^. As expected, the “high dose” micrographs resulted in more visible Thon rings (Figure 2B). Our calculations showed that when high frequencies are present in micrographs and are used for fitting of CTF, defocus can be detected with higher precision (Figure 2C).

**Figure 2.**
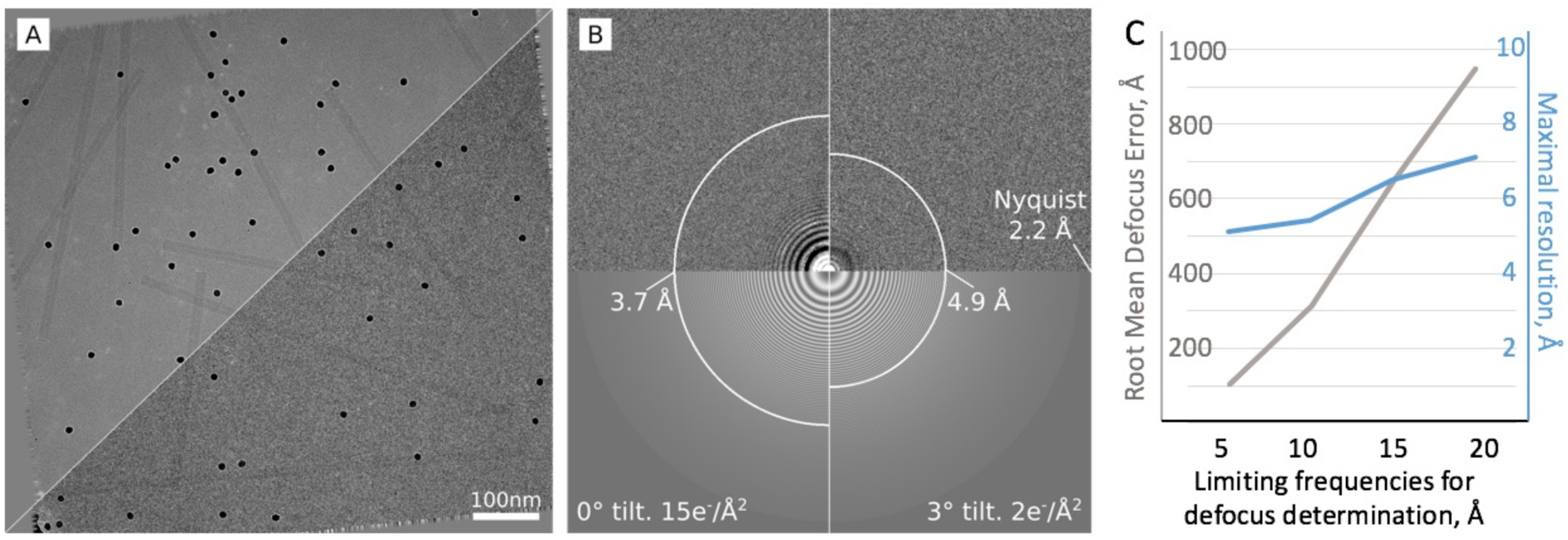
Improved CTF determination for the Hybrid StA. A) A representative micrograph of a 15 e^−^/Å^2^, 0° tilt image (top left) shown together with the corresponding 2 e^−^/Å^2^, 3° tilt image from the same tilt series (bottom right). B) The detected Thon rings for the two images in (A). The white arcs represent the maximum resolution reported by Gctf (Zhang, 2016). C) A quantitative estimation of the relationship between the maximum frequencies used for defocus determination (X-axis) and the defocus error and maximum reported resolution (Y-axis).

We next compared the performance of the conventional StA data processing on hybrid and conventional tomograms of the described TMV datasets. The processing was done using the conventional StA workflow (in Dynamo, see Methods) with the identical parameters and helical symmetry applied. The resulting reconstructions had similar resolutions of 9.8 Å and 10.0 Å respectively (Figure 3A-C). The similarity in resolution for the subtomogram averages resulting from the conventional and hybrid tomograms suggests that collecting data using our hybrid method does not detriment data processing performed by the conventional StA workflow.

**Figure 3.**
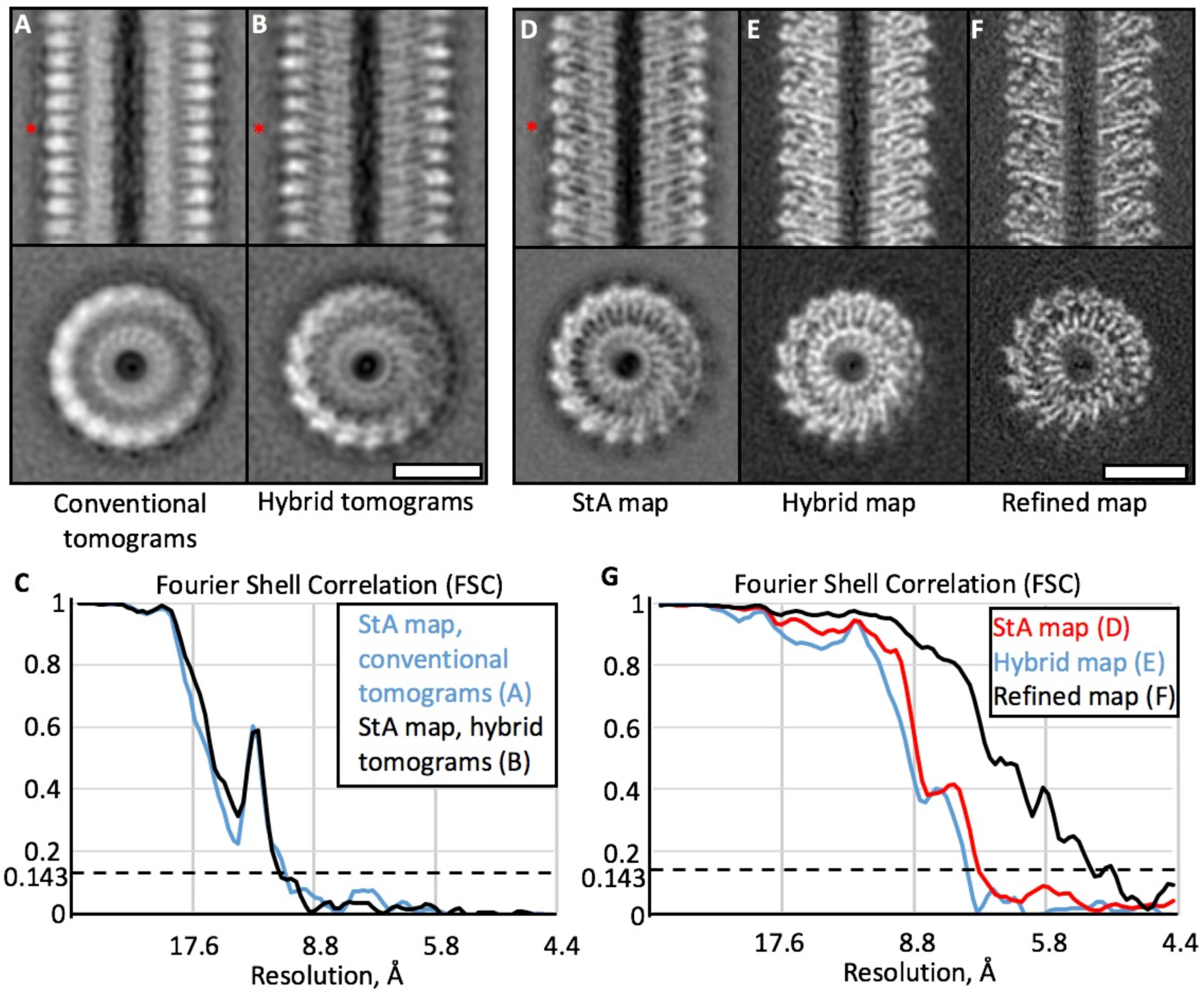
Resolution improvement by the application of *hStA* to TMV. A-C) Structures of TMV (A, B) and the corresponding FSC curves (C) as processed by the conventional subtomogram averaging workflow. The dataset in A was recorded using a conventional dose-symmetric scheme and in B using the proposed hybrid scheme. Note the “halos” observed next to the density in E-F and marked with the asterisks. D-G) Processing of a larger dataset of hybrid tomograms of TMV: (D) conventional StA map; (E) Hybrid map, reconstruction only from the high dose image; (F) and the refined map; (G) the corresponding FSC curves. Scale bars: 10 nm.

Next, using the same settings we recorded a larger dataset of hybrid tomograms of TMV, containing 20,214 asymmetric units. Conventional StA processing with helical symmetry resulted in a 7.2 Å structure (Figure 3D). The Hybrid map, obtained using data only from the “high dose” image, resulted in a 7.5 Å structure (Figure 3E). Interestingly, conventional subtomogram averages had “halos” adjacent to observed protein density (marked with red asterisk in Figure 3A, B, D). To the knowledge of the authors, these “halos” are present to a certain extent in all structures produced by subtomogram averaging. Similar effect has been reported when CTF correction on 2D images has been performed by phase flipping, while the application of an adjusted Wiener filter recovered the original images more precisely (Sindelar & Grigorieff, 2011). Conventional StA processing bases CTF correction on phase flipping (Xiong *et al*, 2009), while in *hStA* Wiener filtration (implemented in Relion) is employed; as a result, while the FSC curves for the conventional StA map and the hybrid map look similar, the hybrid map has a better appearance (Figure 3E). Finally, we used Relion to refine the hybrid map which improved the resolution to 5.2 Å, clearly showing alpha-helical secondary (Figure 3F).

### Structure of an ion channel RyR1 in crowded native membranes

We next applied *hStA* to the structural analysis of ryanodine receptor RyR1 in native SR vesicles purified from fresh rabbit skeletal muscle. We previously reported a structure of RyR1 that reached 12.6 Å from 2,547 particles with C4 symmetry applied (Chen & Kudryashev, 2020) by conventional StA. Here, we recorded a set of hybrid tomograms of the same samples at a higher nominal magnification of 81 000x, corresponding to a pixel size of 1.7 Å (Figure 4A). We picked 2,715 particles and used Dynamo to obtain a StA map of 12.9 Å resolution (Figure 4B), using C4 symmetry. We then exported the particles and performed refinement in Relion applying C4 symmetry. The resulting refined map from 2,563 particles had a resolution of 9.1 Å (Figure 4C, D), and allowed direct observation of secondary structure in the transmembrane and cytoplasmic domains (Figure 4E).

**Figure 4.**
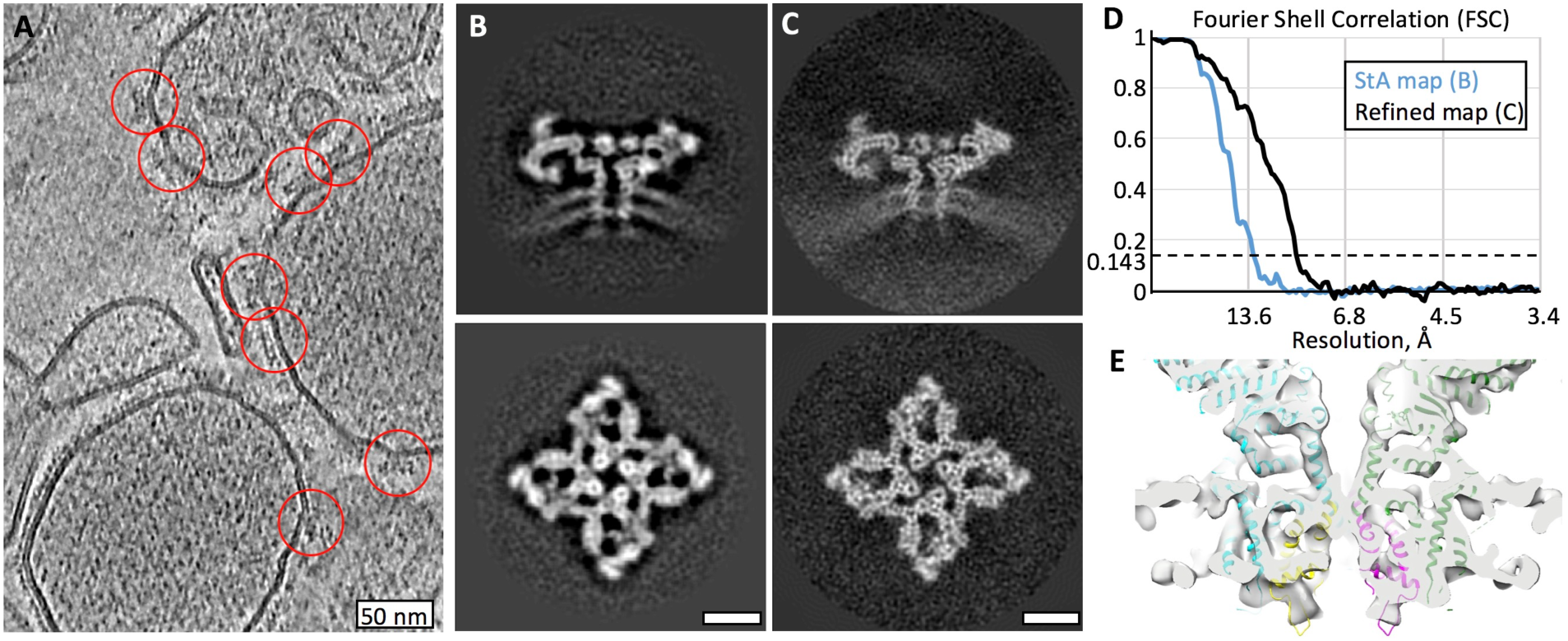
Structure of RyR1 in native membranes at subnanometer resolution. A) A slice through a tomogram of native SR vesicles extracted from rabbit skeletal muscle. Red circles highlight RyR1 particles on the membrane of SR vesicles. Scale bar: 50 nm. B) Slices through a subtomogram average generated from 2,715 particles at resolution of 12.9 Å. C) Slices though a refined map at 9.1 Å resolution generated from 2,563 particles. Scale bars in B, C: 10 nm. D) Corresponding FSC curves between the respective independently processed half sets. E) A volume rendered visualization of the transmembrane domain of the refined map of RyR1 with a rigidly-fit atomic model (PDB ID: *5TB2* from (des Georges *et al*, 2016)).

## DISCUSSION

Here we report a hybrid workflow for subtomogram averaging that redistributes the electron dose across a tilt series, allowing to leverage the image processing tools developed for SPA and StA to achieve significantly higher resolution. We demonstrate that, by using the *hStA* approach, subnanometer-resolution maps can be obtained with data acquired at intermediate magnifications of electron microscopes. We attribute the observed gain in resolution to three main factors:

1. By redistributing more of the available electron dose to the “high-dose” image, the defocus estimation, and therefore the CTF correction can be performed more precisely. Importantly, better CTF correction is performed on the data with higher SNR which is actually used for the final reconstruction.
2. Knowing the height of each particle within its tomogram allows accounting for the defocus gradient within the sample during CTF correction.
3. The ability to compensate for imperfections in tomographic geometry and for beam-induced sample movement in the final refinement step.
4. The use of single-particle-analysis software for reconstruction enables application of Wiener filtering to improve the density quality.

The resolution for the RyR1 dataset improved from 12.9 Å to subnanometer 9.1 Å using 2,563 RyR1 particles and the application of C4 symmetry. The relatively small number of particles was probably the limiting factor for reaching higher resolution. It should be noted that in our examples, we used ∼15% of the total electron dose for the “high-dose” micrograph; however, this fraction or the total electron dose could be increased if the target protein is small or is located in thick ice.

The recently published method *Tygress*, presented by Song and colleagues describes the same idea of recording a higher-dose untilted image and refining the results of StA with SPA tools (Song *et al*, 2020); however, differences exist. First, *Tygress* does not use the untilted high-dose projection for tomogram generation or subtomogram averaging. In our implementation we introduce a normalization procedure which enables all the collected data to be used for subtomogram alignment. Secondly, the implementation details vary: both methods are implemented in MATLAB, in *Tygress*, a *Peet* (Nicastro *et al*, 2006) project is refined with *Frealign* (Grigorieff, 2016) inside Matlab, using a set of configuration files and a GUI to execute the final refinement step. Our approach uses a minimal set of Matlab functions and classes to export a *Dynamo* project into a STAR file for refinement in Relion (Zivanov *et al*, 2018) or other packages. Both implementations provide a significant improvement in resolution: the use of *TYGRESS* allowed obtaining a reconstruction of the outer doublet microtubules from an intact *Tetrahymena thermophila* ciliary axoneme at 10.6 Å from 112,386 particles. Using hStA we reached 9.1 Å using a much lower number of 10,252 asymmetric units; although the differences in the numbers of necessary particles could be partially attributed to the other factors such as ice thickness, differences in hardware.

The optimal scheme for tomographic data collection is still under debate – while the dose-symmetric tilt scheme (Hagen *et al*, 2016) provides better data, it comes at the price of lower throughput (Turoňová *et al*, 2020). In our implementation of hStA only the first “high-dose” image is used for the final reconstruction which relaxes the quality requirements for the remaining projections in the tilt series. Furthermore, hybrid tomographic acquisition and processing may be naturally combined with collecting fast tilt series (Chreifi *et al*, 2019, Eisenstein *et al*, 2019) to increase throughput. If combined with conservatively increasing the angular step and/or reducing the angular coverage of the tilt series, throughput could further increased to one tomogram every few minutes, improving the current throughput by 10-fold. Furthermore, we envisage that *hStA* could be beneficial for fiducial-less samples, such as FIB-milled lamella (Schaffer *et al*, 2019), as the lack of fiducials for tilt series alignment reduces the quality of the resulting tomograms, affecting the quality of the resulting subtomogram averages. Based on our results, we believe that use of hybrid tomograms and application of our *hStA* workflow would be beneficial for a large number of projects aimed at obtaining structures of molecular complexes with subnanometer resolution or higher.

### IMPLEMENTATION AND AVAILABILITY

The modified SerialEM script *HybridDoseSymmetricTomo* is available in Supplementary note 1. The processing workflow is implemented in MATLAB (Mathworks). The test hybrid dataset of TMV has been deposited to EMPIAR along with the alignment parameters for the tomograms, particle locations, and the alignment parameters for the particles in order to produce the hybrid map (EMPIAR-10393). The maps are deposited to EMDB: TMV *hStA* map-EMD –10834, RyR1 *hStA* map – EMD-10840.

The documented scripts are presented in Supplementary note 2, the *dyn2rel* code is deposited to Github (https://github.com/KudryashevLab/dyn2rel). The tomographic tilt series of RyR1 will be released upon publication.

## ACKNOWLEDGEMENTS

We thank Deryck Mills for his expert electron microscopy support and the sample of TMV. We thank Dr. Kendra Leigh for critical reading of the manuscript. The work was funded by the Sofja Kovalevskaja Award from the Alexander von Humboldt Foundation to MK. RS is partially supported by the starter fellowship from SFB807 from the German Research Foundation. YZ is partially supported by the IMPRES international student scholarship from the Max Planck Society. WC is supported by a fellowship from China Scholarship Council. LD is supported by the Max Planck Society.

## AUTHOR CONTRIBUTIONS

YZ modified the data collection script, prepared the TMV cryo samples, collected the data and contributed to data analysis. RS wrote the processing scripts and performed data analysis. LD collected the TMV dataset used for the Figure 3D-G. WC prepared the sample of RyR1, collected data and contributed to data processing for Figure 4. MK designed and supervised the project, contributed to data analysis, acquired funding. MK wrote the manuscript with contributions from RS, YZ, WC, LD.

The authors declare no conflicts of interest.

## METHODS

### Sample preparation and collection of cryo electron tomograms

The TMV sample, 33 mg/mL, was gently mixed with 10 nm protein-A gold nanoparticles (Cell Microscopy Core, Netherlands, UMC Utrecht), with a volume ratio of 1 to 0.8, immediately before plunge-freezing. 3 μL of sample mixed with gold fiducials was then applied on a twice glow discharged (PELCO easiGlow™ Glow Discharge Cleaning System) Quantifoil Au 1.2/1.3 300 mesh grid, blotted using a Mark IV Vitrobot, and plunge-frozen in liquid ethane. Grids were stored in liquid nitrogen until data collection. Tomograms were recorded using two transmission electron microscopes, a first generation FEI Titan Krios and an FEI Titan Krios G2 (Thermo Fisher Scientific) operated at 300 kV, with the specimen maintained at liquid nitrogen temperatures. Tomograms were acquired at a nominal magnification of 64,000x (2.2 Å per pixel in counting mode) using SerialEM with the dose-symmetric script (Hagen *et al*, 2016) and the hybrid variant (Supplementary note 1). Movie stacks with 0.3 e^−^/Å^2^ per frame (1.5-1.8 e^−^ /Å^2^ per projection) were recorded on a post-GIF K2 Summit direct electron detector (Gatan, Inc.) in super-resolution mode. The total electron dose for both the conventional dose-symmetric and hybrid tomograms was kept the same, ∼95 e^−^/Å^2^. The conventional tomograms had the total dose equally distributed over 41 tilt images, from −60 to 60 degrees with a tilting step of 3 degrees. The hybrid tomograms had a zero-tilt image with a total dose of 15-20 e^−^ /Å^2^, with the remaining dose equally distributed over the remaining 40 tilt images over the same angular range.

Sample preparation for RyR1 is described previously (Chen & Kudryashev, 2020), and the same grids prepared for that publication were also used here for imaging on a first generation FEI Titan Krios (Thermo Fisher Scientific) at a nominal magnification of 81,000x corresponding to a pixel size of 1.7 Å per pixel in counting mode on a Gatan K2 Summit electron detector. A total of 47 tomograms of SR vesicles containing RyR1 were collected using the script in Supplementary note 1. An angular range of −60 to 60 degrees was covered with a 3-degree step. The dose for the zero-degree image was 16 e^−^/Å^2^, and 2 e^−^/Å^2^ for the remaining images, with the target defocus set for between 3.5-4.5 μm.

### Image processing

For the conventional processing, each tilt projection was aligned using MotionCor2 (Zheng *et al*, 2017); and the average defocus was determined using Ctffind4 (Rohou & Grigorieff, 2015) or Gctf (Zhang, 2016). The aligned projections were assembled into stacks and aligned using gold fiducials in Imod (Kremer *et al*, 1996), and the resulting aligned tilt stacks were CTF-corrected using *ctfphasefilp* from Imod (Xiong *et al*, 2009). The generated CTF-corrected reconstructions were used for particle picking using the Dynamo catalogue tools (Castaño-Díez *et al*, 2016). Subtomogram averaging was performed using Dynamo (Castaño-Díez *et al*, 2012), versions 1.281 and higher. For TMV independent half-set were generated by assigning different filaments into different sets, these sets were processed independently. For RyR1 independent half-sets were generated by dividing the particles to even and odd.

**For the hybrid datasets**, for the generation of the tilt series all the micrographs were normalized to have the same mean value of 128 and standard deviation of ∼11. The “high dose” images had a mean value of 128 and standard deviation of 4, which was 2.7 times lower, according to the relation (σ_HD_)^2^/ (σ_LD_)^2^ = e^−^_LD_ / e^−^_HD_. The generated stacks were processed according to the conventional StA processing workflow described above. After the conventional processing, the data was exported to Relion 3.0 using a Matlab script described in Supplementary note 2 and a *relion_refine* refinement command was run on two independent half-sets in Relion (Scheres, 2012, Zivanov *et al*, 2018). The description of the *dyn2rel* package, its parameters and an example of usage for the TMV dataset is presented in Supplementary note 2.

For the precision of defocus determination measurements in Figure 2C, 47 high-dose images from the RyR1 dataset were used. Gctf (Zhang, 2016) was used to estimate the defocus and resolution, initially without restricting the frequencies used for the estimation. Then the higher frequencies up to 5, 10, 15 and 20 Å were excluded from calculation using the *–resH* option. The root mean defocus error was calculated by comparing the restricted results to the Gctf result with the full frequency range.

## Supplementary note 1

~~~
MacroName HybridDoseSymmetricTomo
# Adopted from the script of Wim J.H.Hagen (EMBL Heidelberg 2015)
# Roll Buffers A-> H.
# Uses LowDose
# Run eucentric rough and fine
# Track plus K
# Track min L
# Record plus M
# Record min N

########## SETTINGS ##########

step = 3 # stage tilt step in degrees
tilttimes = 10 # multiply by 4 images + 1 image
Tiltbacklash = -3 # negative tilts will be backlashed, must be negative value!

Driftcrit = 3 # Angstrom/second
Driftinterval = 10 # wait time between drift measurements in seconds
Drifttimes = 5 # maximum number of drift measurements before skipping

########## END SETTINGS ##########

ResetClock
echo =====> batchrun HybridDoseSymmetricTomo

tiltangle = 0

CallFunction HybridDoseSymmetricTomo::TiltZero

# prevent runaway focus
AbsoluteFocusLimits -10 10
FocusChangeLimits -2 2

Loop $tilttimes
	# tilt plus1
tiltangle = $tiltangle + $step
CallFunction HybridDoseSymmetricTomo::TiltPlus
	# tilt min1
tiltangle = -1 * $tiltangle
CallFunction HybridDoseSymmetricTomo::TiltMinus
	# tilt min2
tiltangle = $tiltangle – $step
CallFunction HybridDoseSymmetricTomo::TiltMinus
	# tilt plus2
tiltangle = -1 * $tiltangle
CallFunction HybridDoseSymmetricTomo::TiltPlus
EndLoop

TiltTo 0
ResetImageShift
SetDefocus 0
echo =====================================

function TiltZero
# store stage position
ReportStageXYZ
StageX = $ReportedValue1
StageY = $ReportedValue2

# drift and tracking
T
Copy A K
Copy A L
Delay $driftinterval
Loop $drifttimes index
	T
	AlignTo K
		ReportAlignShift
	dx = $reportedValue3
	dy = $reportedValue4
	dist = sqrt $dx * $dx + $dy * $dy
	rate = $dist / $driftinterval * 10
	echo Rate = $rate A/sec
	If $rate < $driftcrit
		echo Drift is low enough after shot $index
		break
	Elseif $index < $drifttimes
		Delay $driftinterval
	Else
		echo Drift never got below $driftcrit: Skipping …
		break
	Endif
EndLoop

# autofocus
G
G
G

# store defocus
ReportDefocus
focusplus = $RepVal1
focusmin = $RepVal1

# acquire the “high-dose” tilt image here
~~~

**Set Exposure R 15**

~~~
R
S
RestoreCameraSet
Copy A M
Copy A N

# store image shifts
ReportImageShift
ISxplus = $RepVal1
ISyplus = $RepVal2
ISxminus = $RepVal1
ISyminus = $RepVal2

## tracking after just to be sure
#T
#Copy A K
#Copy A L
endfunction

function TiltPlus
# tilt stage
TiltTo $tiltangle

# reset stage XY
MoveStageTo $StageX $StageY

# set defocus and image shift
GoToLowDoseArea R
SetDefocus $focusplus
SetImageShift $ISxplus $ISyplus

# drift and tracking
T
AlignTo K
Delay $driftinterval
Loop $drifttimes index
		T
		AlignTo K
		ReportAlignShift
		dx = $reportedValue3
		dy = $reportedValue4
		dist = sqrt $dx * $dx + $dy * $dy
		rate = $dist / $driftinterval * 10
		echo Rate = $rate A/sec
		If $rate < $driftcrit
				echo Drift is low enough after shot $index
				break
		Elseif $index < $drifttimes
				Delay $driftinterval
		Else
				echo Drift never got below $driftcrit: Skipping …
				break
		Endif
EndLoop

# autofocus. Two rounds. Remove one G for single focus round.
G
G

# store defocus
ReportDefocus
focusplus = $RepVal1

# acquire tilt image
R
S

# tracking after
AlignTo M
Copy A M

# store image shifts
ReportImageShift
ISxplus = $RepVal1
ISyplus = $RepVal2

# new track reference
T
Copy A K
endfunction

Function TiltMinus

# tilt stage with backlash
TiltTo $tiltangle
TiltBy $Tiltbacklash
TiltTo $tiltangle

# reset stage XY
MoveStageTo $StageX $StageY

# set defocus and image shift
GoToLowDoseArea R
SetDefocus $focusmin
SetImageShift $ISxminus $ISyminus

# drift and tracking
T
AlignTo L
Delay $driftinterval
Loop $drifttimes index
		T
		AlignTo L
		ReportAlignShift
		dx = $reportedValue3
		dy = $reportedValue4
		dist = sqrt $dx * $dx + $dy * $dy
		rate = $dist / $driftinterval * 10
		echo Rate = $rate A/sec
		If $rate < $driftcrit
				echo Drift is low enough after shot $index
				break
		Elseif $index < $drifttimes
				Delay $driftinterval
		Else
				echo Drift never got below $driftcrit: Skipping …
				break
		Endif
EndLoop

# autofocus. Two rounds. Remove one G for single focus round.
G
G

# store defocus
ReportDefocus
focusmin = $RepVal1

# acquire tilt image
R
S

# tracking after
AlignTo N
Copy A N

# store image shifts
ReportImageShift
ISxminus = $RepVal1
ISyminus = $RepVal2

# new track reference
T
Copy A L
endfunction
~~~

## Supplementary note 2

In this subsection we explain how to export a project from the Dynamo StA project to Relion. For that, purpose we will use the second, larger TMV dataset to recreate the results shown in Figures 3E and Fig 3F. The original stacks, results of the Dynamo project and the necessary alignment files for the use in Imod can be downloaded from the EMPIAR database using the following link: https://www.ebi.ac.uk/pdbe/emdb/empiar/entry/10393/ The files are the final Dynamo table file (.tbl); and the .st, .tlt, .xf and .defocus files for each tomogram. Below we will explain how to use the *dyn2rel* package, and then provide a description of it.

### Example of usage

In the downloaded dataset EMPIAR-10393 we can find all the files needed to export a Dynamo project into a Relion one. To simplify file access, we arranged the tomograms’ files inside folders and named them following a simple naming convention. The downloaded folder should have the following scheme (showing the files for the first two tomograms only):

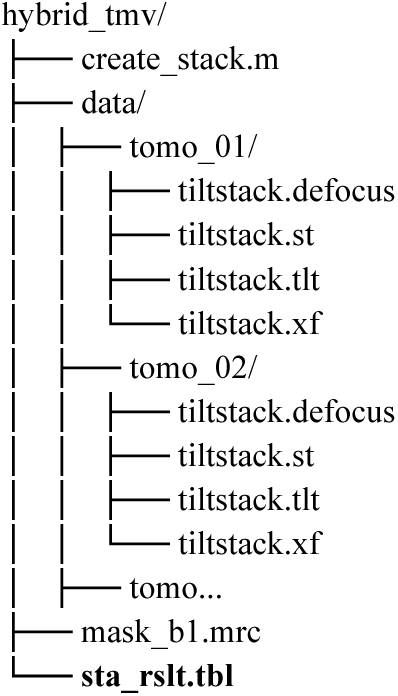

In our case, we used the same name for each .st, .tlt, .ali and .defocus file inside a folder designated for each tomogram. This is not required, however helps to fill the required file name information in the dyn2rel.Tomogram class. The additional three files, “sta_rslt.tbl”, “mask_b1.mrc” and “create_stack.m”, are the resulting table from the Dynamo project, the mask used in said project, and the main exporting script, respectively. The exporting procedure consists of three parts:

1. **Setting up tomograms:** The information of each tomogram is read into a *dyn2rel*.*Tomogram* class, and stored in an array of it. In our case we are using the original unbinned stack, with pixel size of 1.1 Å (recorded in superresolution mode on a K2). We must set up the .st, .xf, .tlt, and .defocus file for each tomogram, along with the full unbinned tomogram size, and the pixel size.
2. **Setting up the exporter:** A *dyn2rel*.*Exporter* is created and configured according to our Dynamo project. In our case, the table was the result of processing a once binned tomogram, so the coordinates and shifts in our table must be multiplied by 2 (exporter.tbl_mul = 2). Also, we want our Relion project to work with a box size of 200 voxels, but binned once too (bin=1). To accomplish this, we set up a cropping size of 400 and a binning of 1 (exporter.out_siz = 400; exporter.out_bin = 1). Finally, we used column 23 in our dynamo table to define to which particle half-set the particles belong, we provide this information for the exporter too (exporter.split_h = 23).
3. **Export the project:** The final step crops the particles from the stacks according to the information on the *dyn2rel*.*Tomogram* array and the table ‘sta_rslt.tbl’ and creates the MRCS stack and the STAR file. In our case, we chose the prefix ‘hyb_b1’ for the project. This creates the ‘hyb_b1.mrcs’ and ‘hyb_b1.star’ files.

The contents of the exporting script are:

% Contents of **create_stack**.**m**

**%% SETTING UP TOMOGRAMS:**

~~~
% Tomogram indices used in **sta_rslt**.**tbl**.
**tomo_ix = [1 3 4 5];**
% as an alternative, we could use:
% tbl = dread(‘sta_rslt.tbl’);
% tomo_ix = unique(tbl(:,20));

% Number of tomograms used in the project:
**N_TOMOS = 4;**
% alternatively we can use
% N_TOMOS = length(tomo_ix);

% Create the dyn2rel.Tomogram array with N_TOMO elements:
**tomos_list = dyn2rel**.**create_tomos_list(N_TOMOS);**

% All the tomograms have the same dimension and pixel size (unbinned).
**tomo_size = [7676, 7420, 1600];**
**pix_size = 1**.**1;**

% Set up the information for each tomogram:
**for i = 1:N_TOMOS**% Set up filenames:
  **base_tomo_dir = sprintf(‘data/tomo_%02d/’**,**i);**
  **stack_name = [base_tomo_dir ‘tiltstack**.**st’];**
  **tlt_name = [base_tomo_dir ‘tiltstack**.**tlt’];**
  **xf_name = [base_tomo_dir ‘tiltstack**.**xf’];**
  **def_name = [base_tomo_dir ‘tiltstack**.**defocus’];**
  
  % Set the information for the i-th tomogram:
  **tomos_list(i)**.**index = tomo_ix(i);**
  **tomos_list(i)**.**set_stack(stack_name**,**pix_size); tomos_list(i)**.**read_ali_files(tlt_name**,**xf_name);**
  **tomos_list(i)**.**set_tomogram_size(tomo_size);**
  **tomos_list(i)**.**set_defocus(def_name);**
**end**
~~~

**%% SETTING UP THE EXPORTER:**

~~~
**exporter = dyn2rel**.**Exporter;**
% In our case, the DYNAMO project was run using binned data (half the size),
% and the stack files (.st) are unbinned. Then, we have to use a factor of 2:
**exporter**.**tbl_mul = 2;**

% The StA map is a binned 200×200×200 volume, to have a hybrid map of the
% same size, we need patches of 400 and then bin the data one time (according
% to the DYNAMO’s dbin command):
**exporter**.**out_siz = 400;**
**exporter**.**out_bin = 1;**

% In our case, we stored the half-set information in the column 23 of the
% table, so we pass that information to the exporter:
**exporter**.**split_h = 23;**

% Finally, we exclude 10% of the particles, the ones with the worst cross
% correlation. Then, we set up xcor_sel to 90% of the particles:
**exporter**.**xcor_sel = 0**.**9;**
~~~

**%% EXPORT THE PROJECT:**

~~~
% Export table **sta_rslt**.**tbl**, using the tomograms in **tomos_list**, By giving
% **‘hyb_b1’** as first argument, the method will create the files **‘hyb_b1**.**star’**
% and **‘hyb_b1**.**mrcs’**, which can be used with RELION:
**exporter**.**exec(‘hyb_b1’**,**tomos_list**,**’sta_rslt**.**tbl’);**
~~~

Once the execution of the script is finished, we reconstruct the hybrid map to obtain the map from Figure 3E

~~~
$ relion_reconstruct –-i hyb_b1.star –-o hyb_b1_rec.mrc --ctf --nr_helical_asu 15 --helical_rise 1.411626 --helical_twist 22.033413 or two half maps:
$ relion_reconstruct –-i hyb_b1.star –-o hyb_b1_h1.mrc --ctf --subset 1 --nr_helical_asu 15 --helical_rise 1.411626 --helical_twist 22.033413
$ relion_reconstruct –-i hyb_b1.star –-o hyb_b1_h2.mrc --ctf --subset 2 --nr_helical_asu 15 --helical_rise 1.411626 --helical_twist 22.033413
~~~

Finally, we refine the results to get the map from Figure 3F

~~~
$ relion_refine –-i hyb_b1.star –-o Refine3D/run_001 --ctf --ref hyb_b1_rec.mrc --angpix 2.2 --o --flatten_solvent --solvent_mask mask_b1.mrc --firstiter_cc --ini_high 8 --offset_range 6 --offset_step 1 --ctf --split_random_halves --auto_refine --norm --scale --pool 4 -- oversampling 1 --healpix_order 6 --sigma_rot 0.1 --sigma_tilt 0.05 --sigma_psi 0.05 --helix --helical_nr_asu 15 --helical_twist_initial 22.0337 --helical_twist_min 21.9 --helical_twist_max 22.1 --helical_twist_inistep 0.05 --helical_rise_initial 1.4123 --helical_rise_min 1.36 -- helical_rise_max 1.46 --helical_rise_inistep 0.1 --helical_z_percentage 0.4 --helical_inner_diameter 20 --helical_outer_diameter 200 -- helical_symmetry_search true --helical_keep_tilt_prior_fixed
~~~

This last command was executed on a computing cluster as a part of a submission script.

Note: For the RyR1 dataset we started with a traditional autorefine Relion project, however, in order to achieve higher resolution, we disabled the autorefinement and enabled the “always_cc” flag. The Relion’s refinement command looks like this:

~~~
$ relion_refine --i autorefine_rslt.star --angpix 3.4 --o Refine3D/cc --sym c4 --iter 20 --flatten_solvent --ref autorefine_rslt.mrc -- solvent_mask mask.mrc --ini_high 15 --offset_range 3 --offset_step 0.2 --ctf --norm --scale --pool 2 --oversampling 1 --healpix_order 7 -- sigma_rot 0.1 --sigma_tilt 0.1--sigma_psi 0.1 --always_cc
~~~

### Description of the dyn2rel package

The Matlab package that perform the exporting procedure is called *dyn2rel*, and can be downloadedfromtheKudryashevlab’sGitHubrepository (https://github.com/KudryashevLab/dyn2rel. To download the package, we go to the desired installation directory and use the *git* command. For this example, we installed the package in the “∼/matlab_packages/dynamo_2_relion/” directory:

~~~
$ cd ∼/matlab_packages/dynamo_2_relion/
$ git clone https://github.com/KudryashevLab/dyn2rel
~~~

To use the package in Matlab we have to add it to its path. In our current Matlab command line, we execute “addpath” and then “help” to validate the correct installation:

~~~
>> addpath ∼/matlab_packages/dynamo_2_relion/
>> help dyn2rel
~~~

If the package was correctly installed, the last command will show the documentation of

*dyn2rel*, which should list the main components of the package:

~~~
Contents of dyn2rel package:

CTF – Holds the CTF’s information.
Exporter – Creates a RELION stack from a DYNAMO tbl and an Array of Tomograms.
Tomogram – Holds a Tomogram’s info: ST, XF, ALI, DEFOCUS files, and its size.
~~~

Here we give a brief description of the main classes and functions of the *dyn2rel* package:

- <u>dyn2rel.CTF:</u> (For internal usage mostly) class that holds the defocus information: U, V, angle, voltage, spherical aberration, amplitude contrast and Bfactor. Additional information can be read using the Matlab command:

~~~
>> help dyn2rel.CTF
~~~
- <u>dyn2rel.Exporter:</u> Class that exports the Dynamo project into Relion for 2D refinement. It reads the Dynamo table, projects the coordinates of each particle into the high-dose projection on the respective stack, crops the corresponding patch, stores it into a MRCS file and writes out a STAR file. The cropping procedure is controlled by the “xcor_sel” property, which sets the percentage of the best particles to be cropped (according to the cross correlation score, column 10 of a Dynamo table). The location of each particle is calculated by adding the shifts (columns 4, 5 and 6) to the position (columns 24, 25 and 26) and then multiplying the result by the “tbl_mul” property. The projection and defocus values are adjusted using the tomogram information (dyn2rel.Tomogram), and the cropped patch size is set by the “out_siz” property. After the patch is cropped, it can be inverted, according to the “invert” property, and binned, according to the “out_bin” property. Additionally, the “split_h” can be used to define how the random half sets are set. Here we give a small summary of the class’ properties grouped by purpose:
  - Particles selection:
    ▪ xcor_sel: fraction of particles to be cropped, according to the cross correlation value (column 10 in a Dynamo table).
  - Table scaling:
    ▪ tbl_mul: Multiplication factor to be applied to the position and shifts on the table to bring the table scaling to the unbinned tomogram scale.
  - Particle’s patch cropping:
    ▪ out_siz: size of the unbinned cropped patch.
    ▪ out_bin: Binning level of the patch. It is applied after the cropping stage, and it is done using Dynamo’s *dbin* command.
    ▪ Invert: Is “true” the values on the patch will be inverted.
  - Extra information:
    ▪ split_h: Sets how the value of “RandomSubset” will be set:
      - split_h < 1: the field will be not set.
      - split_h = 1: even-odd.
      - split_h > 1: the RandomSubset value will be set from the value of the “split_h”-th column in the Dynamo table. Column 23 of Dynamo tables is typically non-assigned.

Finally, the dyn2rel.Exporter class has only one method: exec. This method performs the exporting procedure. It requires 3 input parameters and produces one output:

- dyn2rel.Exporter.exec inputs:
  - out_pfx: (string) base name of the resulting files for the Relion project. The method creates a out_pfx.star file and a out_pfx.mrcs file.
  - tomo_list:(dyn2rel.Tomogram array) contains the information of the tomograms used in the project. It must be created with the dyn2rel.create_tomos_list function.
  - table: (Dynamo table, or filename) table created as a result of a Dynamo project.
- dyn2rel.Exporter.exec output:
  - out_tbl: a subset of table input, containing only the particles that were exported.

For additional information, check the “help” of the class dy2rel.Exporter and its method:

~~~
>> help dyn2rel.Exporter
>> help dyn2rel.Exporter.exec
~~~

- <u>dyn2rel.Tomogram:</u> Class that holds the information of a Tomogram. That information includes unbinned tomogram size (tomo_size), file name of the stack (file_stack), defocus information and the projection information. To set up all the information, the “set_stack”, “read_ali_files”, “set_tomogram_size” and “set_defocus” methods must be used. Additionally, the “index” property must be set, and it is related to column 20 of a Dynamo table. This class also provides the static method “dyn2rel.Tomogram.create_tomos_list”, which must be used to create a list of dyn2rel.Tomogram objects. This list is the one used by the dyn2rel.Exporter class. To see more information, check the help pages of the class:

~~~
>> help dyn2rel.Tomogram
>> help dyn2rel.Tomogram.set_stack
>> help dyn2rel.Tomogram.read_ali_files
>> help dyn2rel.Tomogram.set_tomogram_size
>> help dyn2rel.Tomogram.set_defocus
>> help dyn2rel.Tomogram.create_tomos_list
~~~

